# ActinoMation: a literate programming approach for medium-throughput robotic conjugation of *Streptomyces* spp

**DOI:** 10.1101/2024.12.05.622625

**Authors:** Tenna Alexiadis Møller, Thom Booth, Simon Shaw, Vilhelm Krarup Møller, Rasmus J.N. Frandsen, Tilmann Weber

## Abstract

The genus *Streptomyces* are valuable producers of antibiotics and other pharmaceutically important bioactive compounds. Advances in molecular engineering tools, such as CRISPR, has provided some access to the metabolic potential of *Streptomyces*, but efficient genetic engineering of strains is hindered by laborious and slow manual transformation protocols. In this paper, we present a semi-automated medium-throughput workflow for the introduction of recombinant DNA into *Streptomyces* spp. using the affordable and open-sourced Opentrons (OT-2) robotics platform. To increase the accessibility of the workflow we provide an open-source protocol-creator, ActinoMation. ActinoMation is a literate programming environment using Python in Jupyter Notebook. We validated the method by transforming *Streptomyces coelicolor* (M1152 and M1146), *S. albidoflavus* (J1047), and *S. venezuelae* (DSM40230) with the plasmids pSETGUS and pIJ12551. We demonstrate conjugation efficiencies of 3.33*10^−3^ for M1152 with pSETGUS and pIJ12551; 2.96*10^−3^ for M1146 with pSETGUS and pIJ12551; 1.21*10^−5^ for J1047 with pSETGUS and 4.70*10^−4^ with pIJ12551, and 4.97*10^−2^ for DSM40230 with pSETGUS and 6.13*10^−2^ with pIJ12551 with a false positive rate between 8.33% and 54.54%. Automation of the conjugation workflow improves consistency when handling large sample sizes that facilitates easy reproducibility on a larger scale.

## 1. Introduction

*Streptomyces* have an important role as producers of antibiotics, agricultural agents and enzymes^1–43,4^. Next generation sequencing has increased the number of whole genomes available, revealing the untapped biotechnological potential of these organisms^5–7^. Recent years have also seen the rapid development of molecular genetic tools to engineer *Streptomyces* spp.^8–13^. For example, the advancement of CRISPR tools, such as base-editing^14^, repression^15^, ligation^16^, and recombination^17^ cloning, that have eased the build phase of “Design-Build-Test-Learn” (DBTL)-based metabolic engineering strategies^18^. Despite these advances, engineering efforts are still limited by transformation efficiencies, especially when working with yet uncharacterized strains^19,20^. A widely used method for transforming *Streptomyces* spp. is intergeneric conjugation with *E. coli*^21^. Conjugation involves the co-culture of *Streptomyces* with a conjugative strain of *E*.*coli* containing the plasmid of interest. Although the protocol is conceptually straightforward, conjugation on a large-scale, for example, across a strain collection of thousands of strains, is labor-intensive^10^ and may suffer from low transformation rates^9,22^. Thus, more efficient and scalable workflows for transformation of *Streptomyces* spp. are in high demand.

One method to achieve a higher throughput is with fully, or semi-automated, liquid handling platforms ^23^. These platforms are a robotic pipetting setup that integrates a molecular biology platform to combine synthetic biology and automation engineering to tackle biological problems in high-throughput^23^. Torres et. al. ^24^ divides laboratory robotics equipment into three groups^24^: all-in-one devices, that incorporate the entire experiment into a single setup; multidevices, that can operate both as standalone or combined with other robots for more complex tasks; and, finally, small modular devices, that utilize small modules and can be combined to create expandable systems. All-in-one systems are designed to run the highest number of samples possible on a plug-and-play setup that is easy to learn to use. Many robotic setups are built with large-scale rigid workflows in mind, significantly enhancing industrial applications and delivering very high throughput. However, these setups are expensive and demand a large physical area and require dedicated highly skilled staff to operate. Moreover, not all projects are amenable to rigid automated workflows. Due to lack of standardization of protocols, high capital requirement for setup, and training costs, a classic robotic setup often is not suitable for academic and start-up projects^25,26^.

There are five main bottlenecks in implementing medium-to high-throughput systems (see Figure 1). These bottlenecks have two sides where one is the actual implementation of robotic automation and the other is the way a laboratory currently works which affects the opportunity to automate. Firstly, cost: all-in-one platforms are expensive and dedicated to specific tasks, whereas many research projects necessitate flexible solutions. Hence, the size and the cost of the robotics setups must be relative to the useability requirements and resources of the group. Secondly, programming proficiency: most scientists and laboratory technicians are not sufficiently proficient in both programming and synthetic/molecular biology to automate existing protocols^27^. Thirdly: drain of expert knowledge. Projects can often be treated as “one-and-done,” resulting in highly individualized setups. Insufficient handover of documentation results in a loss of knowledge and capability. This is especially acute in laboratories with high staff turnover. Fourthly, laboratory standardization: poorly documented methodological decisions leads to a lack of experimental standardization, hindering optimization, the ability to compare results across experiments and replication^28–30^. Lastly: blind adherence to established protocols following tips. Often the way a protocol is carried out is affected by the hand-over from a senior user. It could be a tip given by a Postdoc or a note once written on an older version that the current staff now follows. Many lab members also have idiosyncratic adaptations of the same protocol. This mindset impedes the adoption of automation as it can be difficult to decide which version to adapt without thorough evaluation. Together, these challenges collectively impede the widespread use of automation.

**Figure 1.**
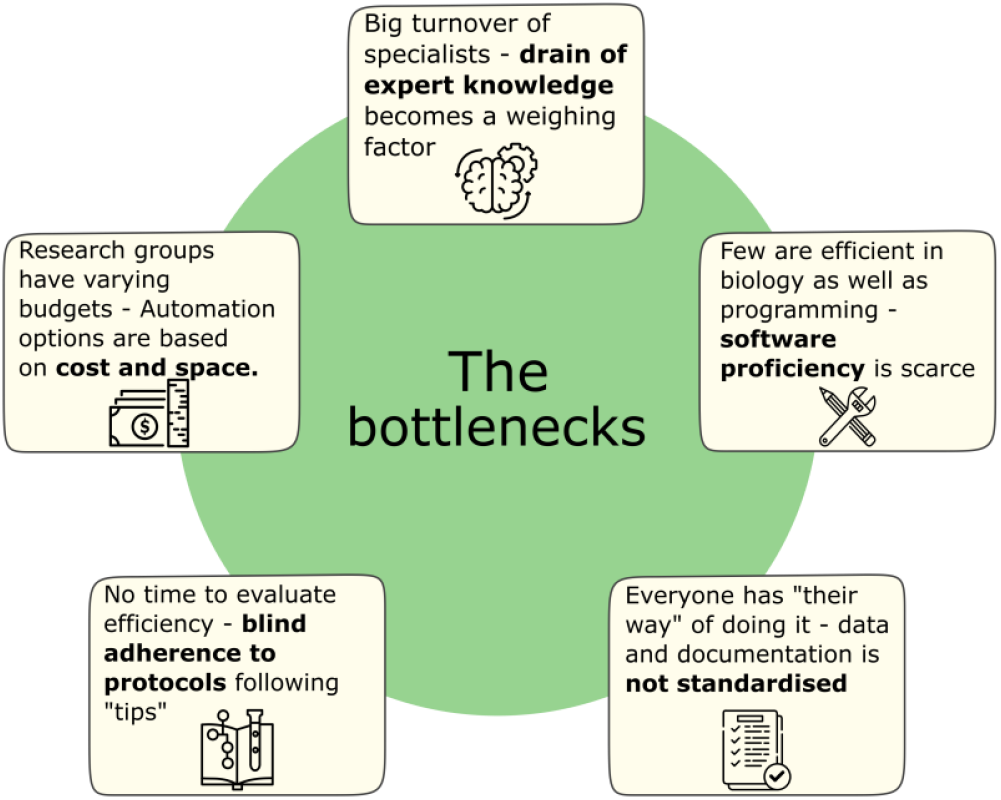
The five bottlenecks to efficiently implementing a medium-to high-throughput workflow in smaller research environments such as academic research groups and start-ups.

Alternatively, modular systems like Opentrons and their OT-2 have gained interest by many academic laboratories due to their affordable price, and flexibility, allowing users to tailor setups to their specific needs, with little requirement of prior knowledge of robotics and programming. Opentrons being powered by Raspberry Pi computers and controlled via Python scripts not only lowers costs but also empowers users to design their own modules, protocols, and cloud-based GUIs. For model bacteria like *E*.*coli* there already exist automatic workflows using Opentrons robots that involves assembly design with open-source Python packages^31,32^ and hardware implementation of already existing cloning platforms^33^. However, setups for handling *Streptomyces* spp. are scarce. Some success has been found using prediction and prioritization of target BGCs via self-resistance genes, cloning and expressing BGCs with high-throughput fermentation and product extraction^34^.

Despite the increased integration of Opentrons workflows^31,35^, papers describing the “design” and “build” of these workflows^23^ are limited. One paper that stands out in this regard is Chitre et. al.^36^ where each step of their setup is detailed and accessible. They encourage other users to replicate and develop on their “open-source hardware” to foster transparency, collaboration and a deeper understanding of equipment functionality while promoting informed design decisions^37^. Furthermore, this openness facilitates cross-disciplinary teamwork, as it allows for colleagues of different educational backgrounds and levels of expertise to come together, therefore mitigating the risk of the knowledge loss bottleneck, should one of the users leave. This creates a shared resource of affordable, customizable setups that other research groups can adopt, saving time and reducing costs.

An increasingly popular method for adapting protocols for robotics is using literate programming to generate scripts for the robot operations. In brief, literate programming leverages recent advances to create computer codes from natural language descriptions. Here among are projects like PyLabRobot^38^, an open-source framework for programming liquid-handling robots and accessories with Python or an large language models (LLM’s) assistant, like GPT-4^39^ and notebook-based tools like Jupyter Notebook^40^ whereby the user interacts with a graphical user interface (GUI) akin to laboratory information management system (LIMS). Literate programming offers non-coders a convenient way to control a robot through a familiar interface without the need for lower-level programming. We believe that the combination of robotics and literate programming can tackle the five bottlenecks we have outlined above so we can take the first steps towards a more accessible medium-throughput approach to the genetic engineering of *Streptomyces* spp..

Here, we present an easily adaptable, cost-effective, and parallelizable workflow to carry out automated interspecies conjugations of *Streptomyces* spp., with a focus on the Design and Build steps of the DBTL cycle. Our workflow encompasses both heat shock transformation of *E. coli* and subsequent conjugation with *Streptomyces sp*. It uses the OT-2 platform and combines it with literate programming of the robotics using Jupyter Notebooks. To validate the system we used it to transform *E. coli* strain Mach1 T1 with two vectors pSETGUS and pIJ12551. We also automated *E. coli to Streptomyces* conjugation in a 96-well format and validated it by conjugating the same plasmids from ET12567/pUZ8002^41,42^ to *S. coelicolor* M1152 and M1146 *S. albidoflavus* J1074 and *S. venezuelae* DSM 40230. Alongside the hardware automation, we created a literate programming protocol-creator implemented in Jupyter Notebooks that utilizes a GUI-like setup in Python. This software assists the user through the preparation of necessary consumables and reagents and creates code for the robotic protocol. Our setup allows for versatile experimental protocol design and execution while reducing the possibility of human errors or execution-bias.

## 2. Material and methods

### 2.1 Microbiological Culture

*E. coli* strain ET12567/pUZ8002^41,42^ and One Shot Mach1 T1 (ThermoFischer; C862003), with and without pSETGUS ^43^ and pIJ12551^44,45^ were cultured in LB media (20 g/L LB broth (Lennox; Sigma-Aldrich; L3022), 10 g/L yeast extract (Thermo Fisher Scientific; LP0021B), 5 g/L sodium chloride (VWR; 470302)) with or without selection (antibiotics at final concentrations; 25 μg/mL chloramphenicol (Sigma-Aldrich; C0378), 50 μg/mL kanamycin sulfate (Sigma-Aldrich; K1377), 100 μg/mL apramycin sulfate (Sigma-Aldrich; A2024), and/or 25 μg/mL nalidixic acid (Sigma-Aldrich; N8878)) as required. For agar plates, the medium was prepared with 2% w/v of agar (Sigma-Aldrich; 05040).

*Streptomyces coelicolor* M1152^19^ and M1146^19^, *S. albidoflavus* J1074^46^ and *S. venezuelae* DSM 40230 were cultured at 30°C^22^ on SFM (soya flour 20 g/L (Fettreduziertes Bio Sojamehl; Hensel, Germany)), D-mannitol (Sigma-Aldrich; M4125)), malt extract 10 g/L (Sigma-Aldrich; 70167), and glucose 4 g/L (Sigma-Aldrich; G7021))) with or without 100 μg/mL apramycin sulfate as required (Sigma-Aldrich; A2024).

### 2.2 Heat shock transformation of chemically competent *E. coli*

The *E. coli* heat shock transformation protocol was carried out according to New England Biolabs^47^. In this method, 50 μL competent cells are thawed on ice, then mixed with 2 μL of plasmid DNA (50 ng, in TE buffer) and chilled for 30 minutes on ice (or at 4°C on the OT-2). The cells are heat shocked for 30 seconds at 42°C, 100 μL room temperature LB media is added, and the sample is sealed and incubated at 37°C for 60 minutes at 250 rpm. Cells are spread with a sterile loop onto warm selection plates (9 cm Petri dish) and incubated overnight at 37°C. To seal the sample a clear adhesive 96-well film (The Applied Biosystems™ MicroAmp™, ThermoFischer) is used.

### 2.3 Conjugation of *Streptomyces* spp

For conjugation, three colonies of *E. coli* ET12467/pUZ8002 with pSETGUS^43^ or pIJ12551^44,45^ from the transformation plate was inoculated into 5 mL LB supplemented with chloramphenicol, kanamycin, and apramycin. The cultures were incubated at 37 °C, with 250rpm orbital shaking for approx. 3 hours before adding it, together with approx. 15 mL LB and antibiotics, to a 50 mL Falcon tube. The cultures, with loosely tightened lids, were incubated at 37 °C, with 250 rpm orbital shaking for approx. 18 hours and grown overnight to reach stationary state (note: unlike the classical protocol^22^, *E. coli* was not grown to a specific OD). The rest of the conjugation followed the protocol described by Kieser^22^ and Gren^48^. After washing and concentrating 10 times, 150 μL of overnight *E. coli* cells were mixed with approximately 10^8^ *Streptomyces* spores in 150 μL 2xYT (tryptone 16 g/L (Millipore; T9410), plated onto SFM agar plates (supplemented to contain a final concentration of 10 mM MgCl_2_ (Sigma-Aldrich; M1028)), and incubated at 30°C for 16-20 hours. The plates were overlaid with 1 mL water containing 0.5 mg nalidixic acid and the plasmid specific selection antibiotic, then potential exconjugants were picked with sterile toothpicks onto SFM agar plates containing nalidixic acid and the antibiotic needed for selection. Conjugation efficiencies were calculated as the number of *Streptomyces* exconjugant CFUs divided by the estimated number of *Streptomyces* CFUs in the conjugation experiment (estimated by serial dilution).

### 2.4 Colony PCR

To screen for successful exconjugants, a pipette tip was used to suspend material from a single colony in 50 μL water in a 96-well PCR plate(NEST, China) and incubated for 20 minutes in a thermocycler (Eppendorf A, Germany) at 95°C. PCR was performed in 10 μL reactions: 10X Dream Taq Buffer (Thermo Scientific; 01186738) 1 μL, 2mM dNTPs [(Thermo Scientific; 2640011) 1 μL, primer pRM4e-apmR-LF (5’-AGCAGCCACTGGTAAC-3’) 10 mM, 0.5 μL, primer pRM4e-apmR-LR (5’-ATGCAGCGTCGTGTT-3’) 10 mM, 0.5 μL, DreamTaq DNA Polymerase (Thermo Scientific; 01188416) 0.025 μL, boiled colony material 1 μL and H_2_0 7 μL. Primers are custom oligos provided by IDT (Integrated DNA Technologies; IA, USA). Thermal cycling (Eppendorf A, Germany) was conducted as follows: initial denaturation at 95°C for 3 minutes, followed by 29 cycles consisting of denaturation at 95°C for 30 seconds, annealing for 30 seconds at 50°C, extension at 72°C for 30 seconds, and a final extension step at 72°C for 5 minutes.

### 2.5 Hardware / Opentron Robotics

The transformation and conjugation protocols created were implemented on the Opentrons OT-2 (Opentrons Labworks Inc., N.Y., USA) using the Opentron API. The specific setup required for each protocol is shown in Figure 2.

**Figure 2.**
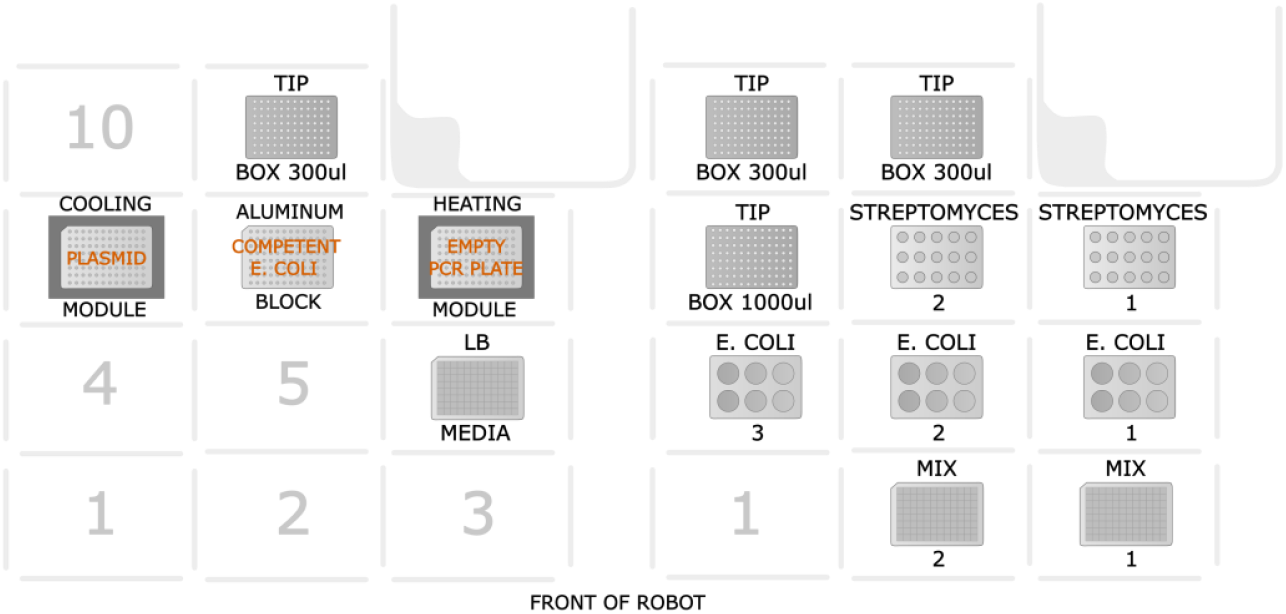
Deck layout for protocols for the transformation with heat shock (left) and conjugation (right) on the OT-2. The numbers indicate the deck slot positions in the robot (notice that position 12 is static and is the trash chute). For the conjugation setup, the numbers on the tip boxes, the tube racks and the plates indicate that the setup allows for multiple plates.

For the heat shock protocol, five positions on the deck are in use. A 96-well deep-well plate is positioned on position 6. A temperature module with an aluminum block and a 96 fully skirted 200 μl PCR plate is placed on position 7 and 9. On position 8 is another aluminum block with a 96 fully skirted 200 μl PCR plate. On position 11 is a 300 μL tip box. For the pipetting head, an 8-channel 300 μL (generation 2) head was used.

For the conjugation protocol consumables are placed depends on the size of the experiment. At maximal capacity ten deck positions are used; on position 2 and 3 a 96-well deep-well plate is positioned. A rack with place for six 50 mL Falcon tubes is placed on positions 4 to 6. A 1000 μL tip box is positioned at position 7. A rack with place for fifteen 15 mL Falcon Tubes is placed on position 8 and 9. Lastly, a 300 μL tip box is placed on position 10 and 11. For the pipetting head, a single-channel 300μL and a single-channel 1000μL head are used.

### 2.6 Jupyter Notebook and Opentrons Software

Protocols and all literate programming manuals were developed in Jupyter Notebook (Python v. 3.9.12). The packages required to run the notebooks are: Opentrons (v. 5.0.2), Zipfile (v. 3.7.0), Ipywidgets (v. 7.6.5), Numpy (v. 1.21.5), Pandas (v. 1.4.2), matplotlib (v. 3.5.1) and Pyisemail (v. 1.4.0).

### The final modified automated workflow is available at

https://github.com/TennaAlexiadisMoeller/ActinoMation and is discussed in full detail in this paper. An in-depth user guide is available at: https://github.com/TennaAlexiadisMoeller/ActinoMation/wiki/How-to-use-guide

## 3. Results and discussion

### 3.1 Workflow design

The goal of this study was to establish a medium-throughput workflow for the transformation of *Streptomyces*. The workflow comprises of a heat shock of chemically competent *E. coli* cells and conjugation with *Streptomyces* on the OT-2 along with some manual steps. This workflow balances throughput with the flexibility required to tailor the experiments for specific projects. There were several general considerations when designing each part of the workflow, including implementation of temperature changes, the positions of hardware modules, loss of cellular material, plastic consumption, plating, precise aspiration and dispensation, and metadata generation.

#### 3.1.1 Heat shock transformation of chemically competent *E*.*coli*

The first step in a conjugation experiment is to transform the plasmid of interest into a conjugative *E. coli* strain, such as *E. coli* ET12467. This is typically achieved via heat shock. Maintaining cells at low temperature before the heat shock is crucial for efficient plasmid uptake. Although storing competent cells on ice is ideal, we did not experience an effect on the cells from the slight temperature difference between 0°C (ice) and 4°C (refrigerator). The Opentrons heating/cooling module operates between 4°C and 95°C, allowing us to adapt the temperature-dependent steps of the protocol, including the 42°C heat shock, to the Opentrons. These hardware modules, however, can only be positioned on the outer edges of the OT-2’s deck (deck positions 1, 3, 4, 6, 7, 9, and 10) due to cord constraints, limiting their number. The competent *E. coli* cells are placed on an aluminium block that has been pre-chilled, wrapped in a bag to prevent condensation, in the freezer. We found that positioning and activation timing significantly impacted performance: The cold module failed to reach the target temperature when placed next to the heating module, or when activated after the heating module, while placing two cold modules together achieved the desired temperatures quickly. Given our findings on placement and temperature stability, we determined that in a single heat shock setup, one 96-well plate of transformants is feasible per robot deck, as shown in Figure 2. Once resolved, we conducted mock wet runs (i.e. using colored water) to test the robot’s aspiration and dispensing accuracy and check for bubble or droplet formation. During testing, we noticed an issue that when using fewer than all eight channels of the multi-channel pipette, the pipetting head would collide with the heating module in one of the first three deck slots of the OT-2. This led us to finalize the deck layout shown in Figure 2. The protocol uses small volumes of competent *E. coli* cells and the *E. coli*-plasmid mixture, and was designed to minimize repeated pipetting and material loss. To ensure minimal cell loss during transfers, an airgap (Figure 3) is used when moving material from the cooling block to the heat shock and then to the deep-well plate with the media. The 8-channel multi-pipette aspirates 100 μl of media from the deep-well plate, then 30 μl of air, and 45 μl of the plasmid and competent *E. coli* mix, leaving some material from the 52 μl to prevent bubbles. Diluting the transformation mix before aspirating minimizes material loss and prevents the formation of bubbles during the aspiration step. Initial issues with droplets and bubbles were resolved by reducing pipetting speed to two-thirds.

**Figure 3.**
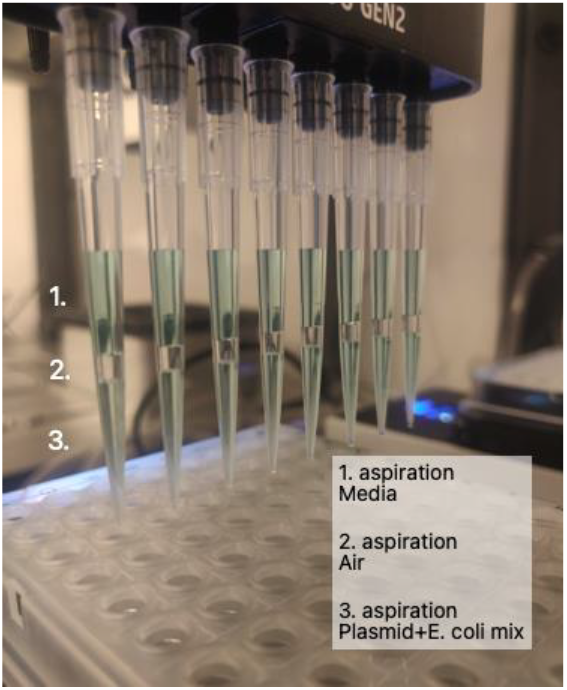
Transfer for the heat shock step in the transformation protocol using an air gap with the in OT-2 multi-channel 300μl pipetting head.

#### 3.1.2 Conjugative transformation of *Streptomyces* with *E*.*coli*

*Streptomyces* conjugation is outlined by Kieser et al.^22^. Briefly, *Streptomyces* spores are incubated with a conjugative *E*.*coli* strain containing the plasmid of interest and a helper plasmid prior to antibiotic selection. When adapting the manual conjugation protocol into a robotic workflow, we designed to minimize the use of consumables and risk of contaminations by optimizing the pipetting patterns. We further modified the Kieser et al. protocol to remove the heat shock of *Streptomyces* spores as we have previously found this step to be unnecessary^48^.

The original protocol calls for a final conjugation mix of 500 μl *E. coli* and *Streptomyces*, whereas we minimized the volumes to 150 μl of each. This allowed us to use a 1000 μl tip for *E. coli*, enabling repeat-dispensing of six wells consecutively. Aspirating extra volume eliminates the need to use the in-built function to blow out the remaining volume and minimizing the risk of bubbles as well as spreading the material for contamination. To reduce the use of tips and reduce unnecessary movement of the pipette head, the tips from the media transfer were reused for the transfer of the *E. coli* strain. To avoid cross-contamination in subsequent steps, new pipette tips were used for the transfer of the *Streptomyces*, which is performed using the single pipetting head with individual 300 μL tips. The original protocol required centrifuging and removing most of the supernatant after mixing of the cultures, before plating. However, with a smaller volume at 300 μL, as opposed to the original protocols 500 μL, we skip this step, saving time and avoiding the need to remove supernatant from a 96-well deep-well plate. In the initial workflow design, we planned to use a 12-well plate for *Streptomyces* and an 8-well plate for *E. coli* to minimize plastic use and streamline the process. However, since *E. coli* are cultured overnight in 50 mL tubes and require two washes, we retained the 50 mL tube format, expanding the capacity from 16 wells to 18 tubes. Also, using 12-well plates for the *Streptomyces* spores limited the combinatorial flexibility. We then considered 15 mL Falcon tubes, which offer several advantages: familiarity for lab colleagues, compatibility with OT-2 racks accommodating up to 15 tubes per deck position, sterile packaging that saves autoclaving time, and screw-top lids that eliminate the need for separate seals. This setup allows for 6 more *Streptomyces* tubes per full setup compared to plates and facilitates sterile transfers between labs. During the wet-run for conjugation, we focused on ensuring proper mixing of experimental strains per the user-specified pattern, with less emphasis on precise pipetting volumes. An initial run checked for droplets that could cause cross-contamination, but with 150 μl in 300 μl tips, no issues with bubbles or aspiration/dispensing were found.

We were not able to automate all steps in the protocol. Chiefly, manual streaking of the conjugation mixture and the addition of the antibiotic overlays is required. Moreover, as satellite colonies often form post-conjugation, we opted not to reduce the surface area of the conjugation plates (i.e. using 48-well plates) and streaked onto standard 9 cm plates. Also, when harvesting spores, we couldn’t grow the *Streptomyces* on small surfaces like 96-well plates, as the protocol requires a significant biomass to produce enough spores for the conjugation experiment.

[We have designed these protocols to adhere with existing lab practices that require high level of flexibility, which in this case led to a more medium-throughput automated robotic workflow. Ultimately, building a high-throughput system requires striking a balance between existing practices and efficiency. Even the most efficient system will fail if users resist change or struggle due to old habits.

#### 3.1.3 Literate programming using Jupyter Notebook

Another objective of our workflow was to standardize data input and output across users, enhancing result comparability. Using the open-source software capabilities of OT-2, we developed fully digital protocols in Python within Jupyter Notebook to automate experiment setup and metadata management. Each workflow was encapsulated in a dedicated, user-friendly notebook designed with literate programming principles, ensuring accessibility for those with minimal or no programming experience. During the design phase, we balanced a sleek interface for user appeal with modularity for component reuse. For the graphical user interface (GUI), we used the iPywidgets framework, enabling interactive data visualization and manipulation without exposing unnecessary code to users. Users are prompted to provide various experiment setup details in the notebook. The notebook then generates the python protocol for the robot, an Excel file listing all necessary materials, and a run-log documenting the robot’s operation, including user identity, run time, and key experiment specifics for future reference. Additionally, it produces visual representations of the final plate layout and images of racks or starting plates with samples (see Figure S1) for traceability, saved alongside in a CSV file. The Opentrons GUI is user-friendly for setup and running the robot but lacks features for a fully automatic and traceable workflow. By developing a system using literate programming in Jupyter Notebook, we eliminated the requirement for individually programming the robot and offered the comfort of using a robotics-protocol with minimal supervision, making it accessible for institutions without access to large biofoundries. Standardized experiment procedures and data management practices mitigate knowledge loss associated with project completion and personnel turnover. Consistent data enables retrospective analysis of experiment runs from a structured perspective, potentially leading to optimized experiment designs. Our automated notebooks provide a simple GUI for users and robust “behind the scenes” data handling. This setup easily customizes the Opentrons protocol, offering greater flexibility, modularity, and integration with data creation, all in one streamlined system.

### 3.2 Workflow implementation

In this section, we outline the detailed workflows for heat shock transformation and conjugation using OT-2 robots. We describe the setup and execution of each protocol, highlighting the use of interactive Jupyter notebooks that guide users through the process, automate critical steps, and ensure consistent results.

#### 3.2.1 Heat shock transformation workflow

The final heat shock transformation workflow entailed the following setup (see Figure 4.a for schematic overview of workflow and Figure 2 for deck layout on the robot): two aluminum blocks are placed by the user on the two temperature modules, located at deck positions 7 and 9, respectively. On the heating module the user adds a fully skirted 200 μL 96-well PCR plate to be heated with the module and at position 11 a box with sterile 200 μL tips placed. The protocol is initiated through the OT-2 graphical user interface (GUI). The two temperature modules need time to reach 4°C and 42 °C. Meanwhile, the user prepares 50 μL plasmid(s) and 2 μL competent *E. coli* cells per conjugation. If low in number this can be done manually, but otherwise the plasmid(s) can be prepared on a fully skirted 200 μL PCR plate using the Mantis (Formulatrix, Dubai) and the competent *E. coli* cells can be prepared on the OT-2 on a fully skirted 200 μL PCR plate. The user also prepares 950 μL per well of warm 37 °C LB media in a 2 mL deep-well 96-well plate. This plate is then placed on the robot at deck position 6, where the plate with plasmid is placed on the cooling module on deck position 7 and the competent cells are placed on a separate freezer-cooled aluminum block on deck position 8. When all material is placed on the robot, the user continues the protocol in the Opentrons GUI. The robot aspirates from the plate with competent cells and dispenses this to the plate with plasmids in the 4°C module, and mixes. This is then incubated for 30 minutes prior to heat shock. Once the robot has completed the run, the user will add a PCR sealing film to the 2 mL deep-well plate and leave it for shaking in a 37°C incubator for 60 minutes at 250 rpm. After incubation, a small volume is plated on the preheated plates and left overnight at 37 °C.

**Figure 4.**
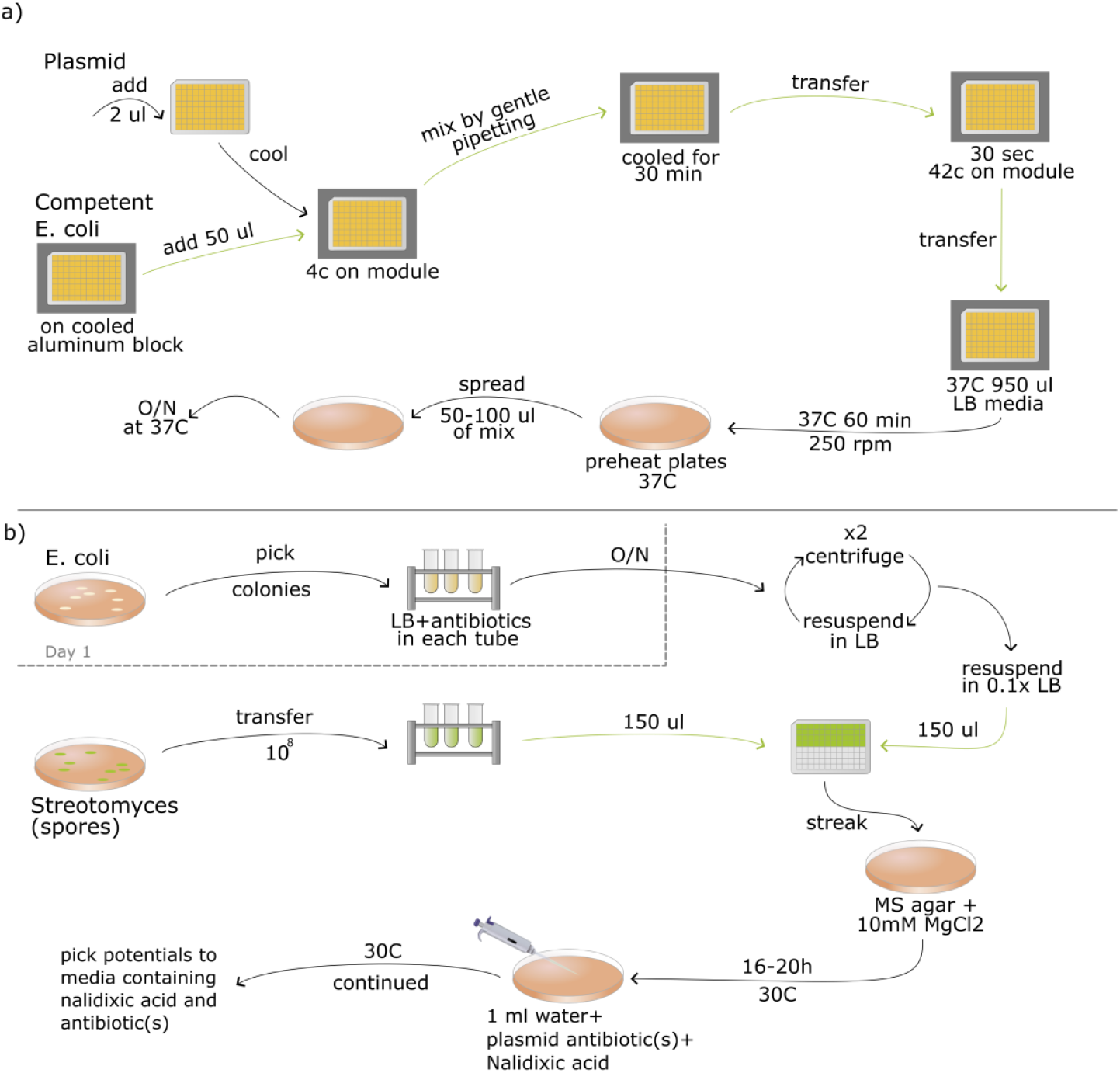
Final automated workflow for E. coli heat shock transformation (a)) and conjugation of Streptomyces spp. (b)) on OT-2. Green arrows indicate steps that are done by the robot.

The conjugation workflow spans three days (see Figure 4.b for schematic overview of workflow and Figure 2 for deck layout on the robot). On the first day, the user picks the 3 colonies from the freshly prepared *E. coli* culture containing transformants from the day before. For this workflow three colonies are added to 5 mL LB media plus antibiotic and then incubated at 250 rpm for 3 hours before it is added to 15 mL LB and antibiotics to a 50 mL Falcon tube with a loose lid and left O/N before starting day two. On the second day, the user will wash the O/N culture two times with equal volume of LB media before resuspending it in 10 percent of the initial volume. The prepared culture is sufficient for 31 individual conjugations reactions. Next, the user will harvest the *Streptomyces* spores into 15 mL Falcon tubes using approx. 5 mL media. Now the user is ready to place everything onto the OT-2 (see Figure 2 for deck layout on the robot). For the conjugation, the user can test up to 30 different *Streptomyces* strains with up to 18 different *E. coli* strains in a single experiment, yielding 192 individual conjugation experiments dispensed in two 96-well deep-well plates. The user places the rack with 15 mL tubes with

*Streptomyces* spores on deck position 9, tubes with the *E. coli* on deck position 6, the 96-well deep-well plates for the conjugation mix on deck position 3 (filled first) and position 2 (filled secondly, only needed if more than 96 combinations). Boxes with 1000 μL tips and 300 μL tips are placed on deck position 7 and 11, respectively. The use can now activate the conjugation protocol via the Opentrons GUI. The OT-2 robot now mixes 150 μL of spores and 150 μL *E*.*coli* into the empty 96-well plates. The 1000 μL tips aspirate and dispense the *E. coli*, while the 300 μL tips aspirate and dispense *Streptomyces* spores. Once the robot has completed the program, the user streaks the content of the wells containing the conjugation mix on MS-agar plates supplemented with 10 mM MgCl_2_ and incubates it O/N at 30 °C. On the final day, an overlay of sterile ddH_2_O with the plasmid specific selection antibiotic(s) and nalidixic acid is added to each plate. The plates are dried in a sterile flow hood and incubated at 30°C until colonies form. See Figure4.b for details.

#### 3.2.3 Jupyter Notebook protocols using literate programming

A complete user guide is available at https://github.com/TennaAlexiadisMoeller/ActinoMation/. The software we have build to create the OT-2 protocols and metadata uses of Jupyter notebooks. These notebooks provide a visual and interactive tool with which users can reproducibly design experiments, and automatically create metadata and Opentrons protocol scripts. There are two separate Jupyter notebooks, one for transformation of *E. coli* and one for conjugation. The software source code is structured to separate those two sections, with a third section (named “common”) which contains code used by both main sections (as seen in Figure S2). For the transformation notebook, users are prompted to upload a tab-separated text file containing the relevant information about the user’s specific antibiotics, plasmids and competent cells. This information is then used throughout the notebook and eventually in the creation of the python script that will control the OT-2. This is illustrated in Figure 5.a.

**Figure 5.**
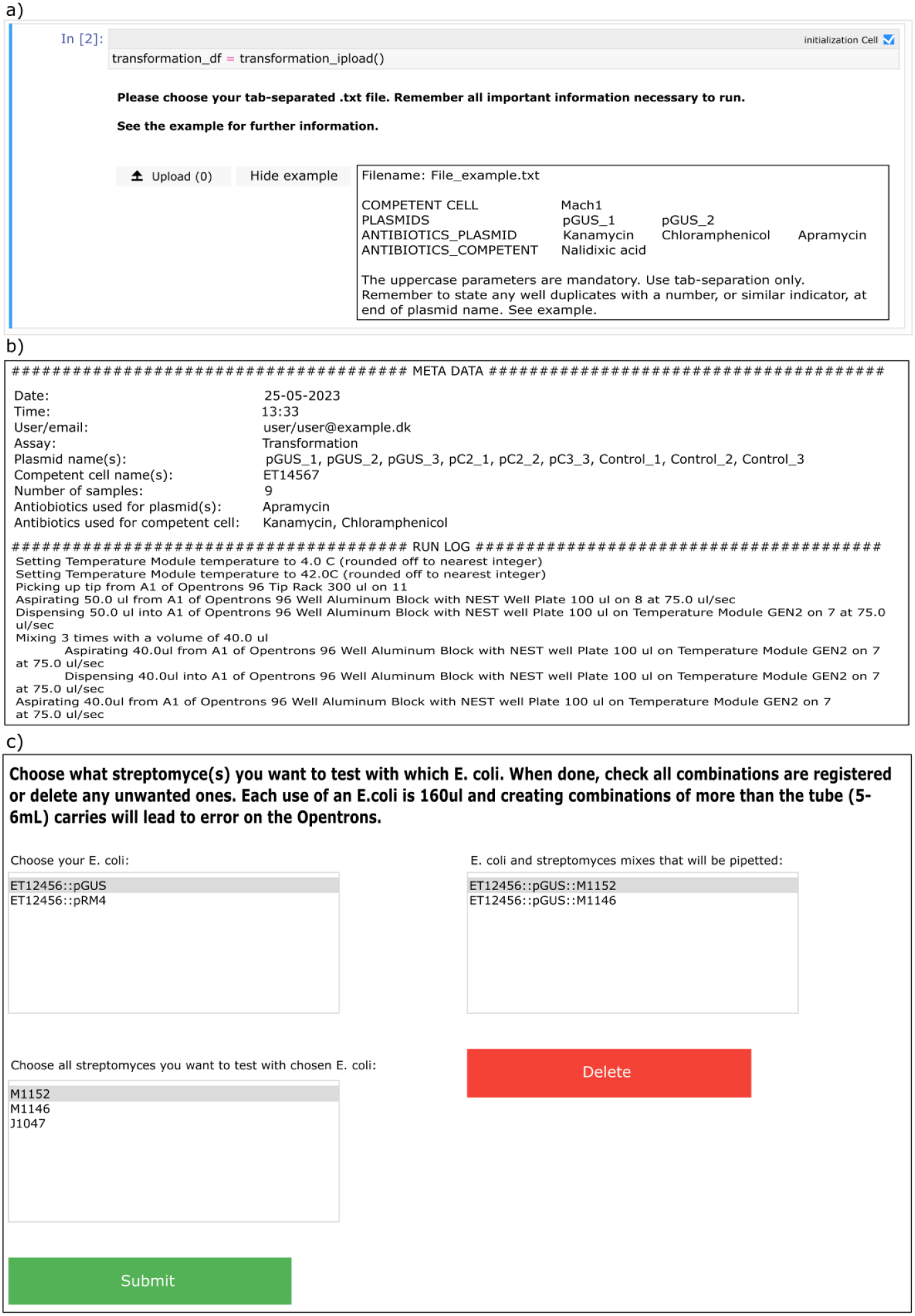
**a)** An example of the guided input process in a notebook. The user can easily view the required input format by clicking on the “Show example” button. **b)** An example of the metadata generated by the notebooks. Including the user provided information along with a step-by-step description of the tasks that the robot will perform. c) The GUI with which a user can easily create their desired mixing combinations, which will be carried out by the robot. The options are derived from the information initially provided by the user.

**Figure 6.**
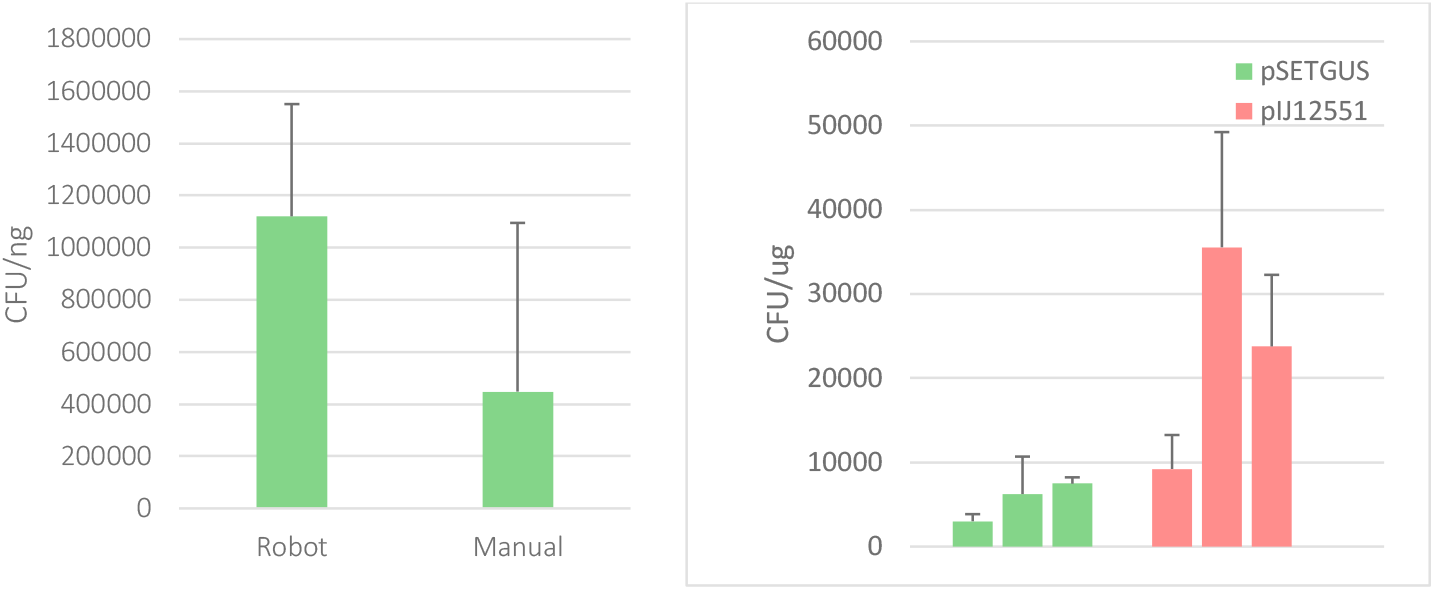
Left: Comparison of efficiencies of transformation of E. coli Mach1 with pSETGUS in experiments with the robot and manual protocol. We included 8 samples for each test, which for the robotic setup equalled to a single column. Right: Transformation efficiency of E. coli ET12567 with pSETGUS and pIJ12551 on the robotic setup. One run included pSETGUS, and pIJ12551. We did three runs with three replicates as well as negative control. Both figures error bars are standard deviations.

As the user progresses through the guided steps of the notebook, a number of outputs will be displayed and saved to disk. These outputs are: a spreadsheet (in XLSX format) containing all necessary consumables, liquids, and materials for future reference; graphical representations of the various plates, and corresponding robot setup, are saved as image files (in PNG format); a text file containing the protocol’s metadata and description (see Figure 5b); and, finally, the generated protocol file that will be uploaded to the OT-2 control system.

A similar process is used in the conjugation notebook. As the conjugation is more complex with mixing of *E. coli* and *Streptomyces*, the notebook GUI allows extra user inputs, which describe the user’s desired mixing pattern (see Figure 5b). The notebook also calculates the minimum amount of LB media necessary for the robot to run without incidence during the overnight culturing with washing. Although ActinoMation was designed specifically for conjugation, these tools can be readily adapted for similar experimental setups. As the Jupyter notebook has a backend of stand-alone modules these can be reused for other workflows and expand the protocol setup and metadata concerning the experimental design in other experiments.

### 3.3 Workflow validation

In this section, we outline the experiments we conducted to validate the two automated robotic workflows and evaluate if they adhere to the expected efficiency of the original manual protocol. Data is presented and highlighted as well as discussed.

#### 3.3.1 E. coli heat shock

To compare the robotic versus the manual protocol we performed transformation experiments with both setups with three independent experiments, each including three replicates. As a proof of concept, we transformed *E. coli* Mach1 T1 (ThermoFischer; C862003) with the plasmid pSETGUS^43^. There was no obvious difference in transformation efficiency between the manual and automated method; the average efficiency of the manual and robotic protocol were 1.12*10^6^ (SD 4.3*10^5^) CFU/ng and 4.48*10^5^ (SD 6.5*10^5^) CFU/ng, respectively. We also compared transformation efficiency to the that of the efficiency stated by the company that supplied the competent cells (Thermo Fischer), which was 1 × 10^9^.

We compared the time to complete each version of the protocol. With the LB media already prepared and ready in the heating cabinet together with the plates the runtime for the experiment was 2 hours and 6 minutes on the robot. With the LB media already prepped and ready in the heating cabinet together with the plates the runtime for the experiment was 2 hours and 18 minutes by hand. Incubation at 37 is the longest step and takes 60 minutes. Therefore, the difference in time won’t be significant in smaller scales. An important consideration when comparing these two methodologies is the impact of prior experience on proficiency. While our familiarity with laboratory procedures instilled confidence when executing the manual protocol, experienced user will be able to prepare the experiment at a faster pace. It’s noteworthy that the primary advantage of utilizing the OT-2 will be realized with a bigger sample size as is shown for the conjugation validation.

#### 3.3.2 Streptomyces conjugation

To test the conjugation workflow, combinations of pSETGUS and pIJ12551 with *S. coelicolor* (M1152 and M1146), *S. albidoflavus* (J1047) or *S. venezuelae* (DSM40230) with 12 replicates of each combination were conducted to fill a full 96-well plate as seen in SI Table 1. Colonies were plated and counted to estimate conjugation efficiency (see Table S2 and Table S3) and three colonies from each experiment was screened using colony PCR (see Figure S3) to estimate the rate of false positives (see Table S4). The conjugation efficiency was calculated as the proportion of ex-conjugant colonies divided by the initial spore concentration. The apparent conjugation efficiency averaged 3.33*10^−3^ for M1152 with pSETGUS and pIJ12551; 2.96*10^−3^ for M1146 with pSETGUS and pIJ12551; 1.21*10^−5^ for J1047 with pSETGUS and 4.70*10^−4^ with pIJ12551; and 4.97*10^−2^ for DSM40230 with pSETGUS and 6.13*10^−2^ with pIJ12551. Being able to screen a higher number of samples per run allows us to evaluate the false positive rate. For M1152 there was a false positive rate 8.33% for both plasmids. For M1146 the rate was 54.54% and 45.45% with pSETGUS and pIJ12551, respectively. J1047 showed no false positives with pIJ12551 but had a rate of 40% with pSETGUS. DSM4023 had a false positive rate of 20% with pSETGUS. With pIJ12551 the rate was 8.33% for DSM4023. Having the false positive rates, the true conjugation efficiency was calculated as seen in table S4. The average true conjugation efficiencies averaged 3.06*10^−3^ for M1152 with pSETGUS and pIJ12551; 1.35*10^−3^ for M1146 with pSETGUS and 1.62*10^−3^ with pIJ12551; 2.02*10^−5^ for J1047 with pSETGUS and 4.70*10^−4^ with pIJ12551; and 4.97*10^−2^ for DSM40230 with pSETGUS and 5.62*10^−2^ with pIJ12551. Interestingly, being able to run a higher number of samples we are here able to see that there might be some strain dependency for conjugation efficiency. While M1152 showed reasonable conjugation efficiency for both plasmids, M1146 were close to equally bad and J1047 might better at accepting pSETGUS than pIJ12551. Unfortunately, conjugation efficiency is not commonly reported which adds to the hindrance of uniformed collecting and presentation of data.

Our aim is to continue developing on both the underlying software and the implementation of robots for a fully automated setup in the future. The introduction of literate programming and modularity in the different steps of a full workflow allows for multiple in the team to work and develop on the setup, even after the current specialists might not be here any longer.

## 4 Conclusion

Our workflow integrates heat shock transformation of *E. coli* and subsequent conjugation with *Streptomyces* spp. using the OT-2 platform, a versatile and cost-effective modular liquid handling system. We enhanced this setup with literate programming via Jupyter Notebooks, providing a user-friendly GUI in Python. Validation involved transforming *E. coli* strain ET12567/pUZ8002 with integrative vectors (pSETGUS and pIJ12551) and conjugating it with 96 strains of *S. coelicolor* M1152 and M1146, *S. albidoflavus* J1074 and *S. venezuelae* DSM 40230.

With a modular workflow like this that focuses on simplicity, we have been able to find an intermediate system between manual and fully-automated biofoundries. This empowers users to design their own modules, protocols, and Python-based GUIs leveraging literate programming and tools like Jupyter Notebook. ActinoMation is freely available, and we hope this will enable other labs to include automation as part of their repertoire and work as a gateway to coding proficiency.

## 5. Acknowledgments

We would like to thank Matiss Maleckis, Kai Blin and Christopher Whitford for guidance in the design of the Opentrons workflow and knowledgeable feedback during testing. We would also like to thank Anny Mais for her assistance in generating test data. This work was funded by grants from the Novo Nordisk Foundation (NNF20CC0035580, NNF22OC0078997).

