## Supporting Information for "ActinoMation: a literate programming approach for medium-throughput robotic conjugation of *Streptomyces* spp"

### SUPPLEMENTAL FIGURES

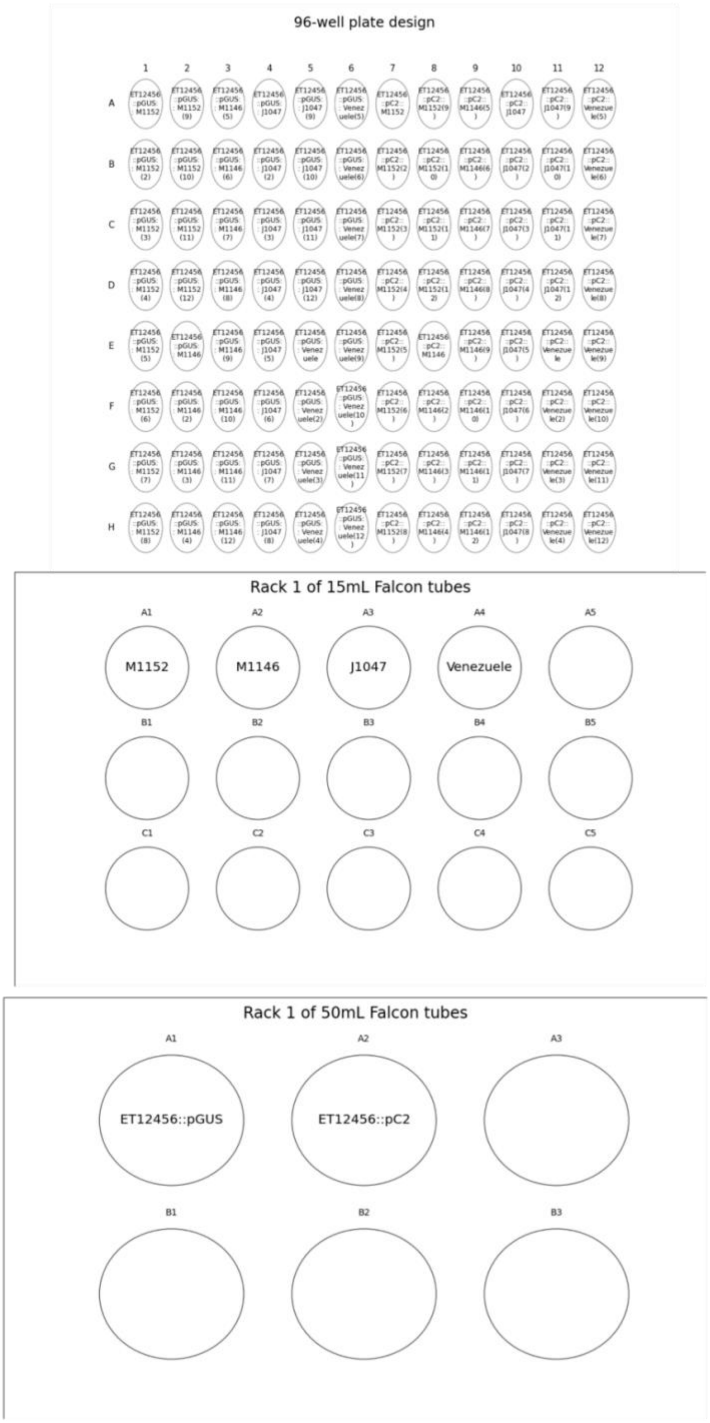

**Figure S1: Example Layouts from ActinoMation.** An example of the visual representation produced by the ActinoMation Jupyter Notebook of the final plate and rack layout including sample names and combinations.

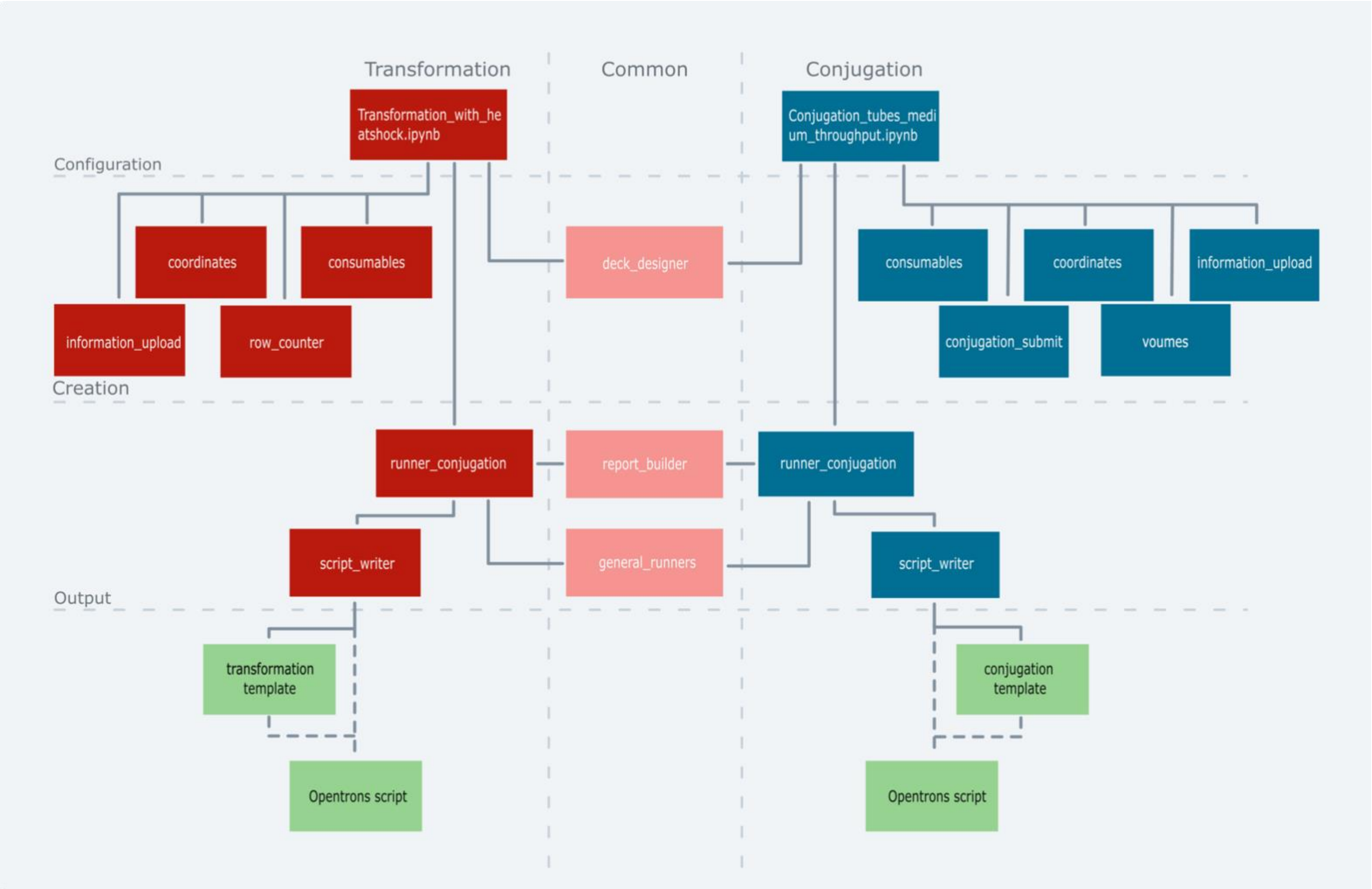

**Figure S2: Dependency tree for the conjugation and transformation workflow notebooks.** Each square represents a .py document containing related functions. The tree is divided into three columns: the left for heat-shock transformation protocol functions, the right for conjugation functions, and the middle for functions shared by both workflows. Horizontally, the tree splits into three sections: "configuration" for user-modifiable protocol functions, "creation" for metadata and Opentrons protocol creation, and "output" for the final product template. The two top squares are Jupyter Notebook files.

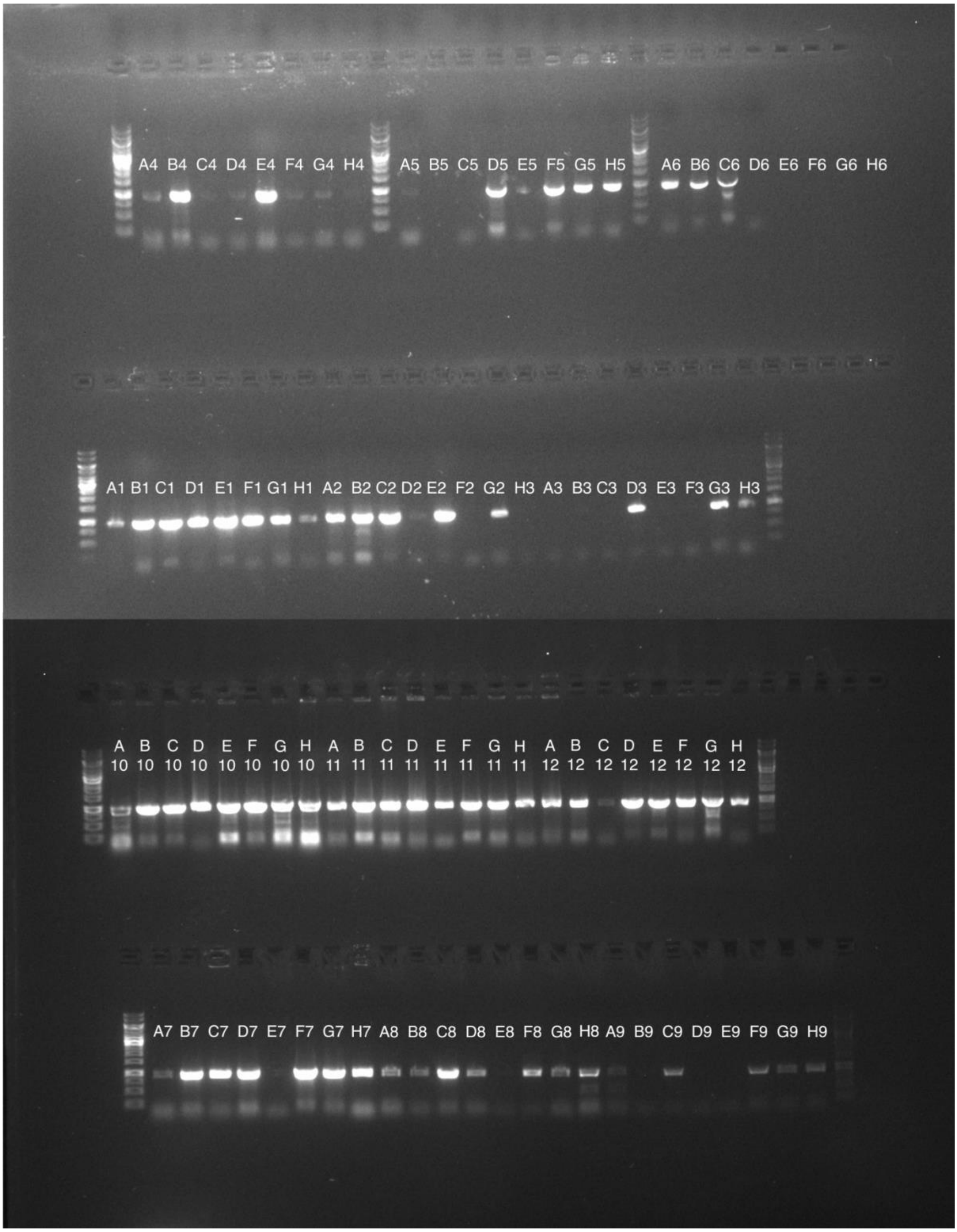

**Figure S3: Agarose gel of colony PCR results.** Conjugation mix of pSETGUS and pIJ12551 with M1152, M1146, J1047 and DSM40230. Letters and numbers correspond with table S1.

### SUPPLEMENTAL TABLES

| | 1 | 2 | 3 | 4 | 5 | 6 | 7 | 8 | 9 | 10 | 11 | 12 |
|---|---|---|---|---|---|---|---|---|---|---|---|---|
| A | ET12456::p<br>SETGUS::<br>M1152 | ET12456::p<br>SETGUS::<br>M1152 | ET12456::p<br>SETGUS::<br>M1146 | ET12456::p<br>SETGUS::<br>J1047 | ET12456::p<br>SETGUS::<br>J1047 | ET12456::p<br>SETGUS::<br>DSM40230 | ET12456::p<br>IJ12551::<br>M1152 | ET12456::p<br>IJ12551::<br>M1152 | ET12456::p<br>IJ12551::<br>M1146 | ET12456::p<br>IJ12551::<br>J1047 | ET12456::p<br>IJ12551::<br>J1047 | ET12456::p<br>IJ12551::<br>DSM40230 |
| B | ET12456::p<br>SETGUS::<br>M1152 | ET12456::p<br>SETGUS::<br>M1152 | ET12456::p<br>SETGUS::<br>M1146 | ET12456::p<br>SETGUS::<br>J1047 | ET12456::p<br>SETGUS::<br>J1047 | ET12456::p<br>SETGUS::<br>DSM40230 | ET12456::p<br>IJ12551::<br>M1152 | ET12456::p<br>IJ12551::<br>M1152 | ET12456::p<br>IJ12551::<br>M1146 | ET12456::p<br>IJ12551::<br>J1047 | ET12456::p<br>IJ12551::<br>J1047 | ET12456::p<br>IJ12551::<br>DSM40230 |
| C | ET12456::p<br>SETGUS::<br>M1152 | ET12456::p<br>SETGUS::<br>M1152 | ET12456::p<br>SETGUS::<br>M1146 | ET12456::p<br>SETGUS::<br>J1047 | ET12456::p<br>SETGUS::<br>J1047 | ET12456::p<br>SETGUS::<br>DSM40230 | ET12456::p<br>IJ12551::<br>M1152 | ET12456::p<br>IJ12551::<br>M1152 | ET12456::p<br>IJ12551::<br>M1146 | ET12456::p<br>IJ12551::<br>J1047 | ET12456::p<br>IJ12551::<br>J1047 | ET12456::p<br>IJ12551::<br>DSM40230 |
| D | ET12456::p<br>SETGUS::<br>M1152 | ET12456::p<br>SETGUS::<br>M1152 | ET12456::p<br>SETGUS::<br>M1146 | ET12456::p<br>SETGUS::<br>J1047 | ET12456::p<br>SETGUS::<br>J1047 | ET12456::p<br>SETGUS::<br>DSM40230 | ET12456::p<br>IJ12551::<br>M1152 | ET12456::p<br>IJ12551::<br>M1152 | ET12456::p<br>IJ12551::<br>M1146 | ET12456::p<br>IJ12551::<br>J1047 | ET12456::p<br>IJ12551::<br>J1047 | ET12456::p<br>IJ12551::<br>DSM40230 |
| E | ET12456::p<br>SETGUS::<br>M1152 | ET12456::p<br>SETGUS::<br>M1146 | ET12456::p<br>SETGUS::<br>M1146 | ET12456::p<br>SETGUS::<br>J1047 | ET12456::p<br>SETGUS::<br>DSM40230 | ET12456::p<br>SETGUS::<br>DSM40230 | ET12456::p<br>IJ12551::<br>M1152 | ET12456::p<br>IJ12551::<br>M1146 | ET12456::p<br>IJ12551::<br>M1146 | ET12456::p<br>IJ12551::<br>J1047 | ET12456::p<br>IJ12551::<br>DSM40230 | ET12456::p<br>IJ12551::<br>DSM40230 |
| F | ET12456::p<br>SETGUS::<br>M1152 | ET12456::p<br>SETGUS::<br>M1146 | ET12456::p<br>SETGUS::<br>M1146 | ET12456::p<br>SETGUS::<br>J1047 | ET12456::p<br>SETGUS::<br>DSM40230 | ET12456::p<br>SETGUS::<br>DSM40230 | ET12456::p<br>IJ12551::<br>M1152 | ET12456::p<br>IJ12551::<br>M1146 | ET12456::p<br>IJ12551::<br>M1146 | ET12456::p<br>IJ12551::<br>J1047 | ET12456::p<br>IJ12551::<br>DSM40230 | ET12456::p<br>IJ12551::<br>DSM40230 |
| G | ET12456::p<br>SETGUS::<br>M1152 | ET12456::p<br>SETGUS::<br>M1146 | ET12456::p<br>SETGUS::<br>M1146 | ET12456::p<br>SETGUS::<br>J1047 | ET12456::p<br>SETGUS::<br>DSM40230 | ET12456::p<br>SETGUS::<br>DSM40230 | ET12456::p<br>IJ12551::<br>M1152 | ET12456::p<br>IJ12551::<br>M1146 | ET12456::p<br>IJ12551::<br>M1146 | ET12456::p<br>IJ12551::<br>J1047 | ET12456::p<br>IJ12551::<br>DSM40230 | ET12456::p<br>IJ12551::<br>DSM40230 |
| H | ET12456::p<br>SETGUS::<br>M1152 | ET12456::p<br>SETGUS::<br>M1146 | ET12456::p<br>SETGUS::<br>M1146 | ET12456::p<br>SETGUS::<br>J1047 | ET12456::p<br>SETGUS::<br>DSM40230 | ET12456::p<br>SETGUS::<br>DSM40230 | ET12456::p<br>IJ12551::<br>M1152 | ET12456::p<br>IJ12551::<br>M1146 | ET12456::p<br>IJ12551::<br>M1146 | ET12456::p<br>IJ12551::<br>J1047 | ET12456::p<br>IJ12551::<br>DSM40230 | ET12456::p<br>IJ12551::<br>DSM40230 |

**Table S1: Plate design for conjugation on the Opentrons 2.** Each cell contains the names of the strains usided in combinations of ET12456::pSETGUS or ET12456::PIJ12551 with M1152, M1146, J1047 and DSM40230.

| | 1 | 2 | 3 | 4 | 5 | 6 | 7 | 8 | 9 | 10 | 11 | 12 |
|---|---|---|---|---|---|---|---|---|---|---|---|---|
| **A** | >100 | >100 | >100 | 0 | 3 | 71 | >100 | >100 | >100 | 63 | 91 | >100 |
| **B** | >100 | >100 | >100 | 5 | 0 | 58 | >100 | >100 | >100 | 55 | 76 | >100 |
| **C** | >100 | >100 | >100 | 5 | 0 | NA | >100 | >100 | >100 | 52 | 69 | >100 |
| **D** | >100 | >100 | >100 | 5 | 8 | NA | >100 | >100 | >100 | 13 | >100 | >100 |
| **E** | >100 | >100 | >100 | 4 | 57 | NA | >100 | >100 | >100 | >100 | >100 | >100 |
| **F** | >100 | >100 | >100 | 5 | 113 | NA | >100 | >100 | >100 | >100 | >100 | >100 |
| **G** | >100 | >100 | >100 | 3 | 74 | NA | >100 | >100 | >100 | >100 | >100 | >100 |
| **H** | >100 | NA | >100 | 1 | NA | NA | >100 | >100 | >100 | 85 | 4 | >100 |

**Table S2: Exconjugant colonies counted on MS agar plates.** Each cell corresponds to the conjugation experiment described in Table S1. Cells with NA are shown were data was not counted due to aspiration errors.

| | 1 | 2 | 3 | 4 | 5 | 6 | 7 | 8 | 9 | 10 | 11 | 12 |
|---|---|---|---|---|---|---|---|---|---|---|---|---|
| A | 0.03333333 | 0.03333333 | 0.00296296 | 0 | 1.1215E-05 | 0.04733333 | 0.03333333 | 0.03333333 | 0.00296296 | 0.00039252 | 0.00056698 | 0.06666667 |
| B | 0.03333333 | 0.03333333 | 0.00296296 | 1.8692E-05 | 0 | 0.03866667 | 0.03333333 | 0.03333333 | 0.00296296 | 0.00034268 | 0.00047352 | 0.06666667 |
| C | 0.03333333 | 0.03333333 | 0.00296296 | 1.8692E-05 | 0 | NA | 0.03333333 | 0.03333333 | 0.00296296 | 0.00032399 | 0.00042991 | 0.06666667 |
| D | 0.03333333 | 0.03333333 | 0.00296296 | 1.8692E-05 | 2.9906E-05 | NA | 0.03333333 | 0.03333333 | 0.00296296 | 8.0997E-05 | 0.00062305 | 0.06666667 |
| E | 0.03333333 | 0.00296296 | 0.00296296 | 1.4953E-05 | 0.038 | NA | 0.03333333 | 0.00296296 | 0.00296296 | 0.00062305 | 0.06666667 | 0.06666667 |
| F | 0.03333333 | 0.00296296 | 0.00296296 | 1.8692E-05 | 0.07533333 | NA | 0.03333333 | 0.00296296 | 0.00296296 | 0.00062305 | 0.06666667 | 0.06666667 |
| G | 0.03333333 | 0.00296296 | 0.00296296 | 1.1215E-05 | 0.04933333 | NA | 0.03333333 | 0.00296296 | 0.00296296 | 0.00062305 | 0.06666667 | 0.06666667 |
| H | 0.03333333 | NA | 0.00296296 | 3.7383E-06 | NA | NA | 0.03333333 | 0.00296296 | 0.00296296 | 0.0005296 | 0.00266667 | 0.06666667 |

**Table S3: Conjugation efficiency.** The conjugation efficiency calculated as  the final number of ex-conjugants from Table S2 divided by the initial spore concentration.

| | 1 | 2 | 3 | 4 | 5 | 6 | 7 | 8 | 9 | 10 | 11 | 12 |
|---|---|---|---|---|---|---|---|---|---|---|---|---|
| A | 0.00305666 | 0.00305666 | 0.00134696 | 0 | 1.8692E-05 | 0.04733333 | 0.00305666 | 0.00305666 | 0.00161629 | 0.00039252 | 0.00056698 | 0.06111334 |
| B | 0.00305666 | 0.00305666 | 0.00134696 | 3.1153E-05 | 0 | 0.03866667 | 0.00305666 | 0.00305666 | 0.00161629 | 0.00034268 | 0.00047352 | 0.06111334 |
| C | 0.00305666 | 0.00305666 | 0.00134696 | 3.1153E-05 | 0 | NA | 0.00305666 | 0.00305666 | 0.00161629 | 0.00032399 | 0.00042991 | 0.06111334 |
| D | 0.00305666 | 0.00305666 | 0.00134696 | 3.1153E-05 | 4.9844E-05 | NA | 0.00305666 | 0.00305666 | 0.00161629 | 8.0997E-05 | 0.00062305 | 0.06111334 |
| E | 0.00305666 | 0.00134696 | 0.00134696 | 2.4922E-05 | 0.038 | NA | 0.00305666 | 0.00161629 | 0.00161629 | 0.00062305 | 0.06111334 | 0.06111334 |
| F | 0.00305666 | 0.00134696 | 0.00134696 | 3.1153E-05 | 0.07533333 | NA | 0.00305666 | 0.00161629 | 0.00161629 | 0.00062305 | 0.06111334 | 0.06111334 |
| G | 0.00305666 | 0.00134696 | 0.00134696 | 1.8692E-05 | 0.04933333 | NA | 0.00305666 | 0.00161629 | 0.00161629 | 0.00062305 | 0.06111334 | 0.06111334 |
| H | 0.00305666 | NA | 0.00134696 | 6.2305E-06 | NA | NA | 0.00305666 | 0.00161629 | 0.00161629 | 0.0005296 | 0.00244454 | 0.06111334 |

**Table S4: True conjugation efficiency.** The efficiency is calculated on data from table S3 and the false positive rates calculated in the main text.
